# GABAergic modulation of conflict adaptation and response inhibition

**DOI:** 10.1101/2022.03.03.482762

**Authors:** Eduardo A. Aponte, Kaja Faßbender, Jakob Heinzle, Ulrich Ettinger

## Abstract

Adaptive behavior is only possible by stopping stereotypical actions to generate new plans according to internal goals. It is response inhibition —the ability to stop actions automatically triggered by exogenous cues— that allows for the flexible interplay between bottom-up, stimulus driven behaviors, and top-down strategies. In addition to response inhibition, cognitive control draws on conflict adaptation, the facilitation of top-down actions following high conflict situations. It is currently unclear whether and how response inhibition and conflict adaptation depend on GABAergic signaling, the main inhibitory neurotransmitter in the human brain. Here, we applied a recently developed computational model (SERIA) to data from two studies (N=150 & 50) of healthy volunteers performing Simon and antisaccade tasks. One of these datasets was acquired under placebo-controlled pharmacological enhancement of GABAergic transmission (lorazepam, an allosteric modulator of the GABA-A receptor). Our model-based results suggest that enhanced GABA-A signaling boosts conflict adaptation but impairs response inhibition. More generally, our computational approach establishes a unified account of response inhibition and conflict adaptation in the Simon and antisaccade tasks and provides a novel tool for quantifying specific aspects of cognitive control and their modulation by pharmacology or disease.

**Author Summary:** Our capacity to prepare for situations that afford conflicting responses (conflict adaptation) and to stop our immediate impulses in these scenarios (response inhibition) are the hallmark of cognitive control. As these abilities require both the *stopping* or *slowing* of response tendencies, a natural question is whether they are mediated by inhibitory neurotransmission in the brain. Here, we combined computational modeling with two experiments to investigate how conflict adaptation and response inhibition interact with each other (experiment 1) and how these are modulated by lorazepam (experiment 2), a positive modulator of the GABA-A receptor, one of the main inhibitory receptors in the human brain. Using our computational model to disentangle conflict adaptation and response inhibition, our results indicate that while lorazepam impaired response inhibition, it improved conflict adaptation. Thus, our results suggests that conflict adaptation is mediated by GABA-A neurotransmission.

## Introduction

When confronted with sudden changes in circumstances, cognitive control becomes imperative: this not only involves stopping the previous course of action, but also selecting and executing a new plan tailored to the changing environment (1). In the motor domain, actions are often programmed and executed automatically - for example, turning the eyes to an unexpected visual stimulus - and need to be stopped in time so that an alternative action can be executed. When environmental cues induce such response conflicts, one major challenge is to inhibit reflexive or prepotent behaviors that start without reflection or planning, i.e. exercise *response inhibition* (2, 3). Response inhibition is one component of cognitive control in anticipation of upcoming challenges (4). Additionally, cognitive control manifests through the facilitation of strategic or goal-directed actions and typically increases following the experience of response conflicts, a phenomenon called *conflict adaptation* (4–6).

Despite our considerable understanding of the neurobiology of response inhibition (7–11) and conflict adaptation (4,12–15), it remains unclear to which extent stopping automatic behaviors in favor of goal-directed actions relies on GABAergic signaling, the main inhibitory neurotransmitter in the brain. It is also unknown whether the behavioral adjustments that follow high conflict situations are mediated by GABAergic signaling. However, indirect evidence indicates that response inhibition is associated with GABAergic neurotransmission (16–20).

One limitation of previous studies that tried to pin down the role of GABAergic signaling in response inhibition and conflict adaptation is the absence of computational models that statistically formalize the interplay between controlled and automatic behaviors. Indeed, while dual-process models postulate the existence of controlled and automatic actions (21–23), mathematical formalizations of their interaction are still rare (but see (24, 25)).

Recently, we introduced the *Stochastic Early Response, Inhibition and late Action* (SERIA) model for the antisaccade task (Aponte et al., 2017), one of the main paradigms used to measure response inhibition when secondary actions are required (see Fig. 1). SERIA combines the “horse-race” model, used in the stop signal task (27, 28), with a linear ballistic accumulation model that decides between controlled actions. The interplay between automatic and controlled behaviors is mediated by a latent inhibitory process that races against the automatic, fast process. In a series of studies (26,29,30), we have shown that SERIA accurately explains reaction time (RT) distributions and error rates (ER), and predicts several features of the antisaccade task in a variety of conditions.

**Figure 1:**
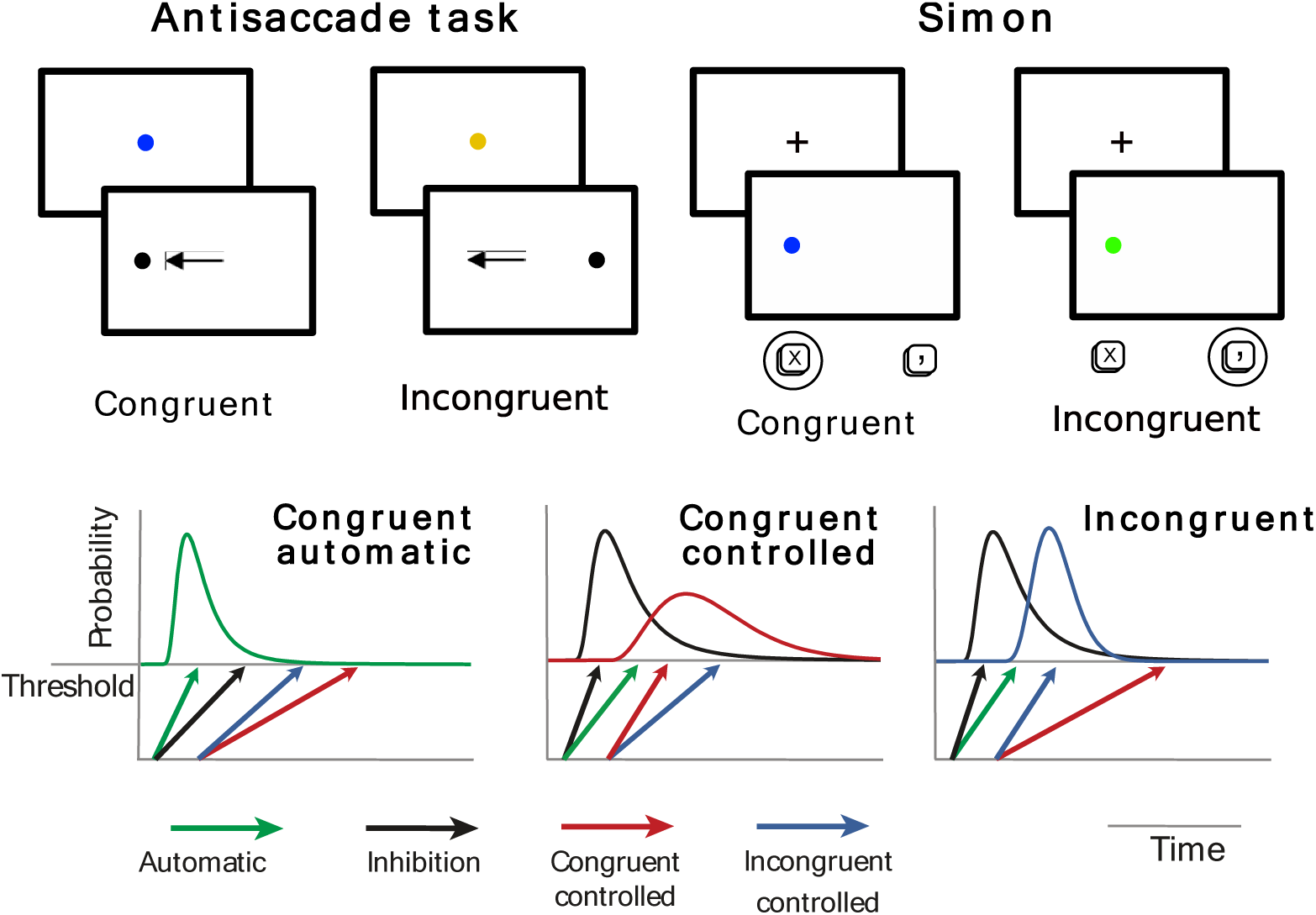
Antisaccade task. A central fixation cue was presented for 1000 to 2000ms. Its color (blue or yellow) indicated subjects to saccade to the peripheral stimulus (congruent trial) or to saccade in the opposite direction (incongruent trial). The peripheral stimulus was presented for 1000ms. **Simon task.** Subjects were instructed to press a left (‘x’) or right (‘,’) key depending on the color (blue or green) of a peripheral cue (display duration 1500ms) following a fixation period of 500ms. On congruent trials, the right-left location of the cue and the correct button matched each other; on incongruent trials, they were in opposite locations. **SERIA model.** Reaction times and actions are assumed to be the outcome of a race to threshold between independent linear accumulators (processes), whose slopes take different, random values on each trial. Initially, the automatic process starts after a short delay from the cue presentation. This process can be stopped by the inhibitory process, if the latter is the first to hit threshold. When this occurs, the outcome of the race between two controlled processes that represent congruent and incongruent responses decides the action. The reaction time on a trial is assumed to be the threshold-hit-time of the corresponding process.

A second paradigm commonly used to measure response inhibition and conflict-induced cognitive control is the Simon task (31). On incongruent trials, an irrelevant spatial feature of the stimulus conflicts with the cued response (for instance, a green colored cue on the right indicating a left key press, when only the color of the cue is relevant). This condition is characterized by slower and more error-prone actions compared to congruent trials, in which the response and the irrelevant spatial location of the stimulus coincide. The difference in RT and ER between incongruent and congruent trials is called the Simon effect, or, more generally, the congruency effect. The advantage of congruent trials reverses after incongruent trials (i.e., high conflict trials), as conflict induced control slows congruent responses and facilitates incongruent responses (4, 32).

In this study, we set out to investigate the role of GABA signaling in response inhibition and conflict adaptation during the antisaccade and Simon tasks. In the first experiment, we confirmed the validity of SERIA in the antisaccade task and demonstrated that the model can be extended to the Simon task. Specifically, SERIA offers a quantitative and comprehensive explanation of the Simon effect, its time course, and its reversal after high conflict trials. To this effect, data from 164 healthy adults performing both tasks were collected and SERIA was applied to trial-by-trial RT.

In the second experiment, we applied the same computational model to a recently published dataset (33) in order to determine the effects of a pro-GABAergic drug on response inhibition and conflict adaptation. Fifty subjects performed the same protocol as in Exp. 1 under placebo, 0.5 and 1.0 mg of the benzodiazepine lorazepam, an unspecific, positive allosteric modulator of the GABA-A receptor. In addition to the expected sedative effect of lorazepam, we found that enhanced GABA-A signaling *boosted* conflict adaptation but *impaired* response inhibition.

## Results

The primary goal of Exp. 1 was to verify that response inhibition and conflict adaptation can be explained using SERIA as a single, unified probabilistic model of the antisaccade and Simon tasks. This entails showing that SERIA captures the main qualitative *and* quantitative features of subjects’ responses. Thus, we examined if the mean reaction times (RT) and error rates (ER) as well as the RT distributions could be predicted by our model after fitting trial-by-trial responses. Note that the model was never exposed to descriptive statistics (e.g., mean reaction times) or their distributions. Rather, the single input to the model was the list of actions (congruent or incongruent responses) and the corresponding RT of every subject. Therefore, we used the generative nature of the model to *predict* mean statistics and the shape of the RT distributions based on the distribution of the posterior parameters.

The second goal of Experiment 1 was to dissect the explanation that SERIA offers for the congruency effect and for conflict adaptation. Hence, we investigated how different model-parameters change across congruent and incongruent trials, and how these changes interact with the conflict level of previous trials.

### Experiment 1

In Exp. 1, from 164 subjects, two were excluded from the analysis of the Simon task because of the elevated number of errors or missing trials (see Methods). The number of excluded subjects increased t o 12 in the antisaccade task, as 11 subjects had 50% or more trials excluded and one subject’s ER was higher than 80%.

For clarity, in the following, “conflict level” refers to the *N-1* trial. Congruent trials are considered low conflict trials, and incongruent trials high conflict trials.

Fig. 2 displays the mean RT and ER in all conditions. As expected, in the antisaccade task, subjects were slower (Δ = 77ms; *P* < 10^―5^) and generated more errors (Δ = 24%; *P* < 10^―5^) on incongruent trials compared to congruent trials. No conflict adaptation was evident in this task, as the congruency effect *in*creased slightly after high conflict trials (8ms; P=0.023). Regarding ER, we again found no evidence of conflict adaptation as the congruency effect was significantly *higher* after high conflict trials compared to low conflict trials (Δ = 9%; *P* = 0.004).

**Figure 2:**
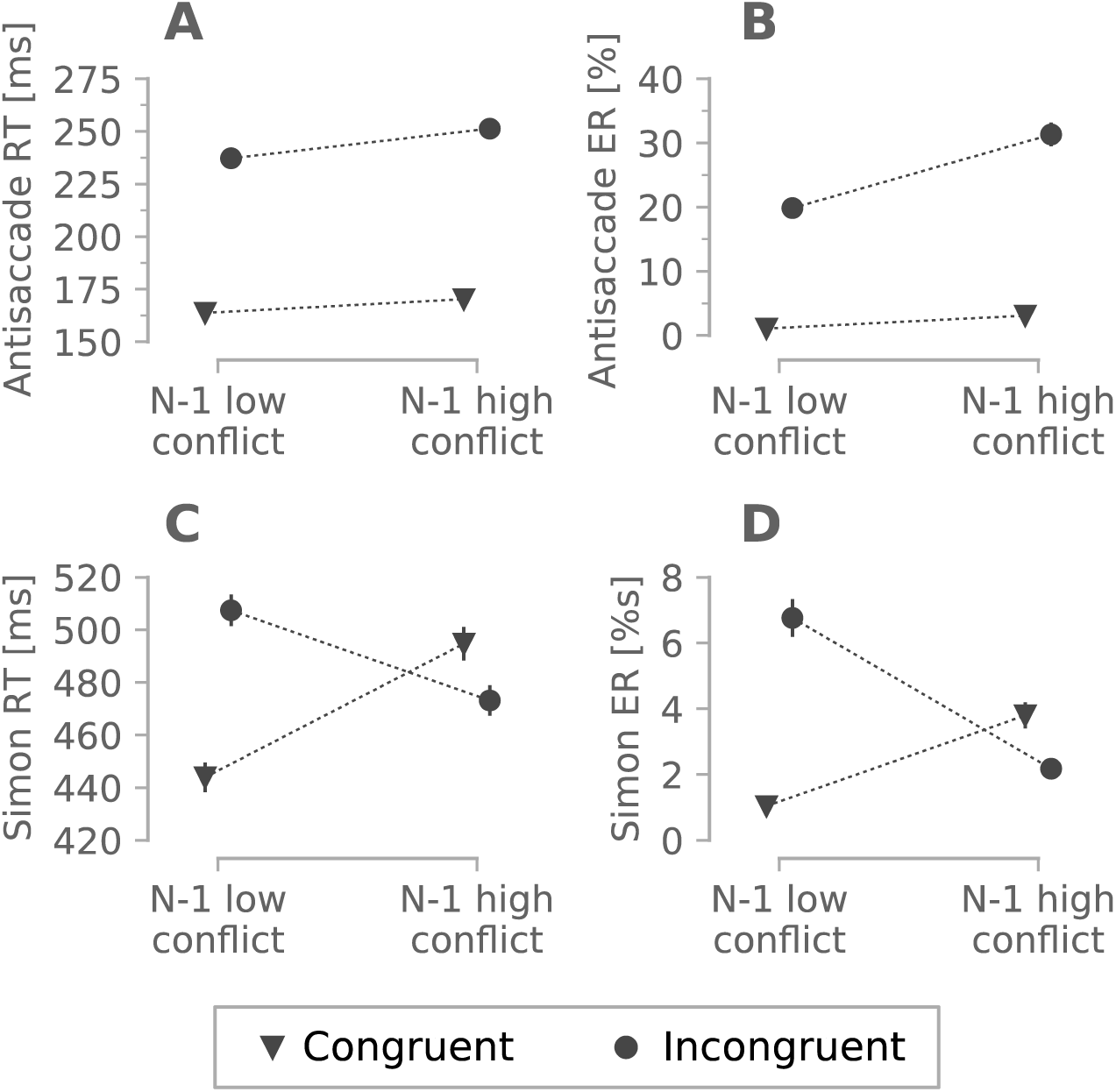
**A)** Mean RT in the antisaccade task. **B)** Mean ER in the antisaccade task. C) Mean RT in the Simon Task. **D)** Mean ER in the Simon task. N-1 refers to the corresponding previous trials. Error bars display the sem.

In the Simon task (Fig. 2C&D), incongruent responses were slower (Δ = 21*ms*,*P* < 10^―5^) and more error prone than congruent responses (Δ = 2%,*P* < 10^―5^). More importantly and in contrast to the antisaccade task, conflict adaptation was observed following high conflict trials. Indeed, the congruency or Simon effect was 63ms after low conflict trials and -22ms after high conflict trials, resulting in a significant interaction between conflict and congruency (Δ = 85*ms*,*P* < 10^―5^). This interaction was driven by two different factors: congruent responses were slower after high conflict trials (Δ = 51*ms*;*P* < 10^―5^) whereas incongruent responses were faster (Δ = ―34*ms* ;*P* < 10^―5^). ER followed the same pattern (see Fig. 2D), with a inversion — from positive to negative — of the Simon effect indicated by a significant interaction (*P* < 10^―5^).

Could SERIA capture the RT distributions of congruent and incongruent trials in both tasks? To answer this question, we fitted SERIA to the RT of each subject in each condition. All group fits displayed here are the weighted average over subjects.

Fig. 3A&D demonstrate that all distributions were fitted with great accuracy. Moreover, the predicted and empirical mean RT and ER closely matched each other (Supp. Fig. 1.) A possible objection here is that congruent responses are not bimodally distributed and consequently there is no evidence that these were generated by two different processes. To answer this objection, we compared SERIA to a simpler model in which all congruent responses originate from a single unimodal distribution. Bayesian model comparison clearly demonstrated that the explanatory power of models (balance between fit and complexity) profited from including a controlled process in the generation of congruent responses (see Supp. Table 1 & 2), even after controlling for the number of parameters. This replicates and extends our previous findings (26, 30).

**Figure 3:**
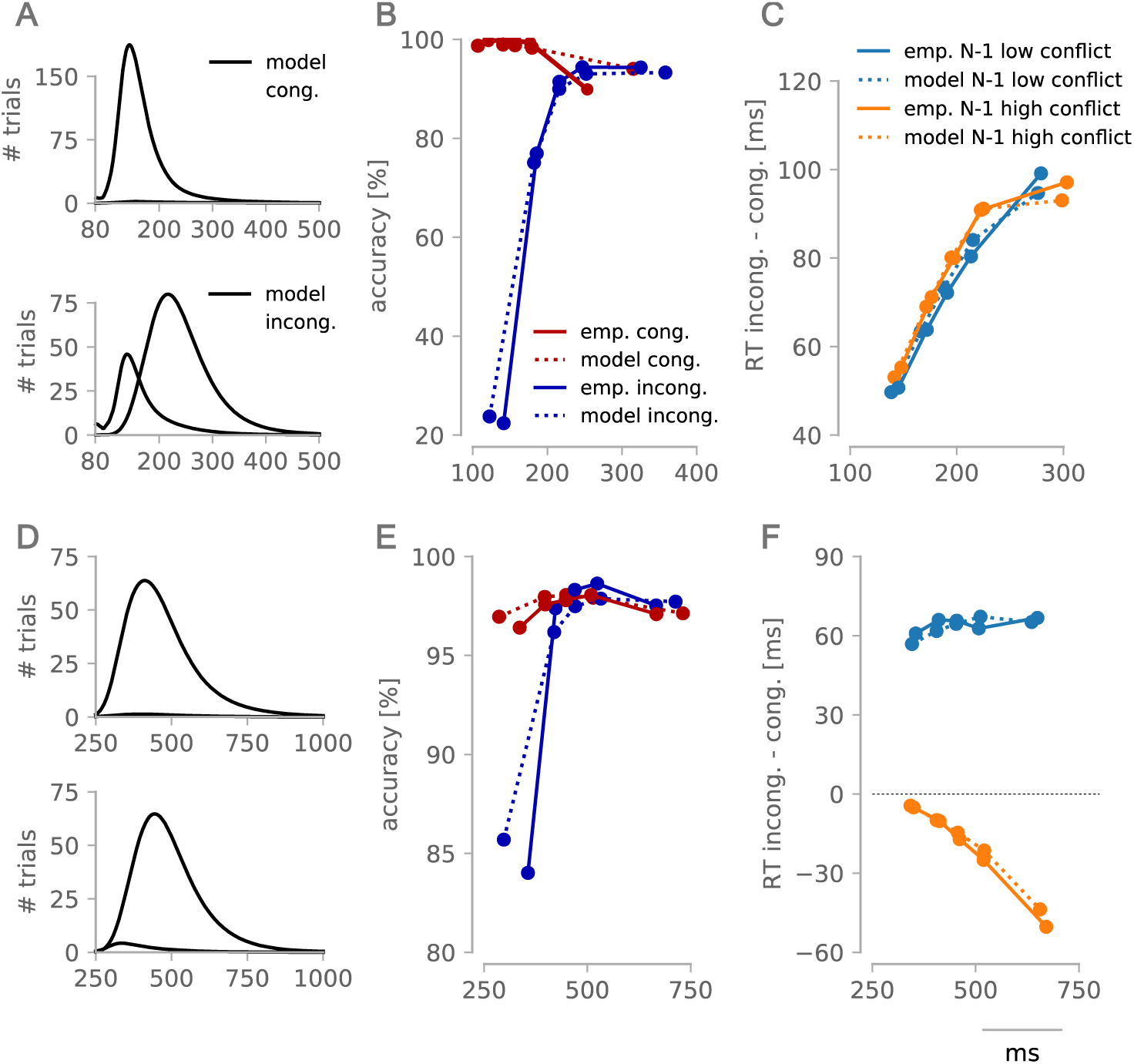
Model fits and distributional analysis. Top row: Antisaccade task. Bottom row: Simon task. **A)** Reaction time (RT) distributions in the antisaccade task. Histograms show empirical data and solid black lines display the model fits. **B)** Accuracy function in the antisaccade task. Accuracy plots were generated by sorting trials in RT percentiles (20, 40, 60, 80 and 100%) and plotting the mean accuracy in each percentile against the corresponding mean RT. **C)** Delta plot in the antisaccade task. As with the accuracy plots, in delta plots trials are binned in RT percentiles and the congruency effect (the difference between mean RT on incongruent and congruent trials) is plotted against the pooled mean RT. **D-F)** RT distribution, accuracy and delta plot in the Simon task, similar to A-C.

Rather than merely predicting the RT histograms, we aimed to reproduce and explain the time course of the congruency effect revealed by delta plots (21). In this analysis, trials are binned in RT quantiles and either the accuracy or the RT congruency effect (i.e., the difference in RT between incongruent and congruent trials) are plotted against the quantile-specific mean RT.

As shown in Fig. 3B&E, the accuracy function followed the same pattern in both tasks, with errors on incongruent trials occurring predominately at low latencies, suggesting that most errors are indeed inhibition failures. Yet, errors were still possible even at the highest time bins, with similar error rate in congruent and incongruent trials. In general, SERIA could accurately reproduce the accuracy functions.

Despite other qualitative similarities between the two tasks, the Simon effect and the antisaccade cost for RT followed widely different time courses (see Fig. 3C&F). The antisaccade RT cost was always positive and increased with latency in all conditions. By contrast, after high conflict trials, the Simon effect was negative and declined as a function of latency. Yet, after low conflict trials, the Simon effect was positive and changed only minimally across time bins. Qualitatively, the model captured the delta plots with great accuracy in both tasks.

Having shown that indeed SERIA can correctly capture response inhibition and conflict adaptation in the Simon task, we can now ask how the model explains the negative Simon effect after high conflict trials, and why is the slope of the delta plot negative in this condition?

To answer the first question, we note that according to SERIA congruent responses can be generated by either the controlled or the automatic process. Thus, the congruency effect can be approximated by the weighted difference between the RT of incongruent responses and the two types of congruent responses (controlled and automatic):

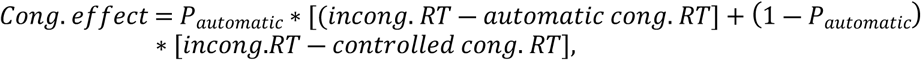

where *P_automatic_* is the probability of an automatic response. This probability weights the contribution of the automatic (congruent) process to the overall mean congruent response RT. To understand the congruency effect and how conflict-induced adaptation interacted with it, we examined the contribution of controlled and automatic responses separately.

In the Simon task, after low conflict trials, there was only a small RT difference between controlled congruent and incongruent responses (Δ = 9ms; Fig. 4D). Despite this small difference, the contribution of fast, automatic responses (approximately 30% of congruent responses, Fig. 4F) led to an overall large Simon effect (predicted by the model to be Δ = 60ms) as automatic responses (Fig. 4E) were on average much faster than controlled incongruent responses (Δ = 155ms). In other words, after low conflict trials, the bulk of the Simon effect was caused by the difference between controlled incongruent and automatic congruent responses.

**Figure 4:**
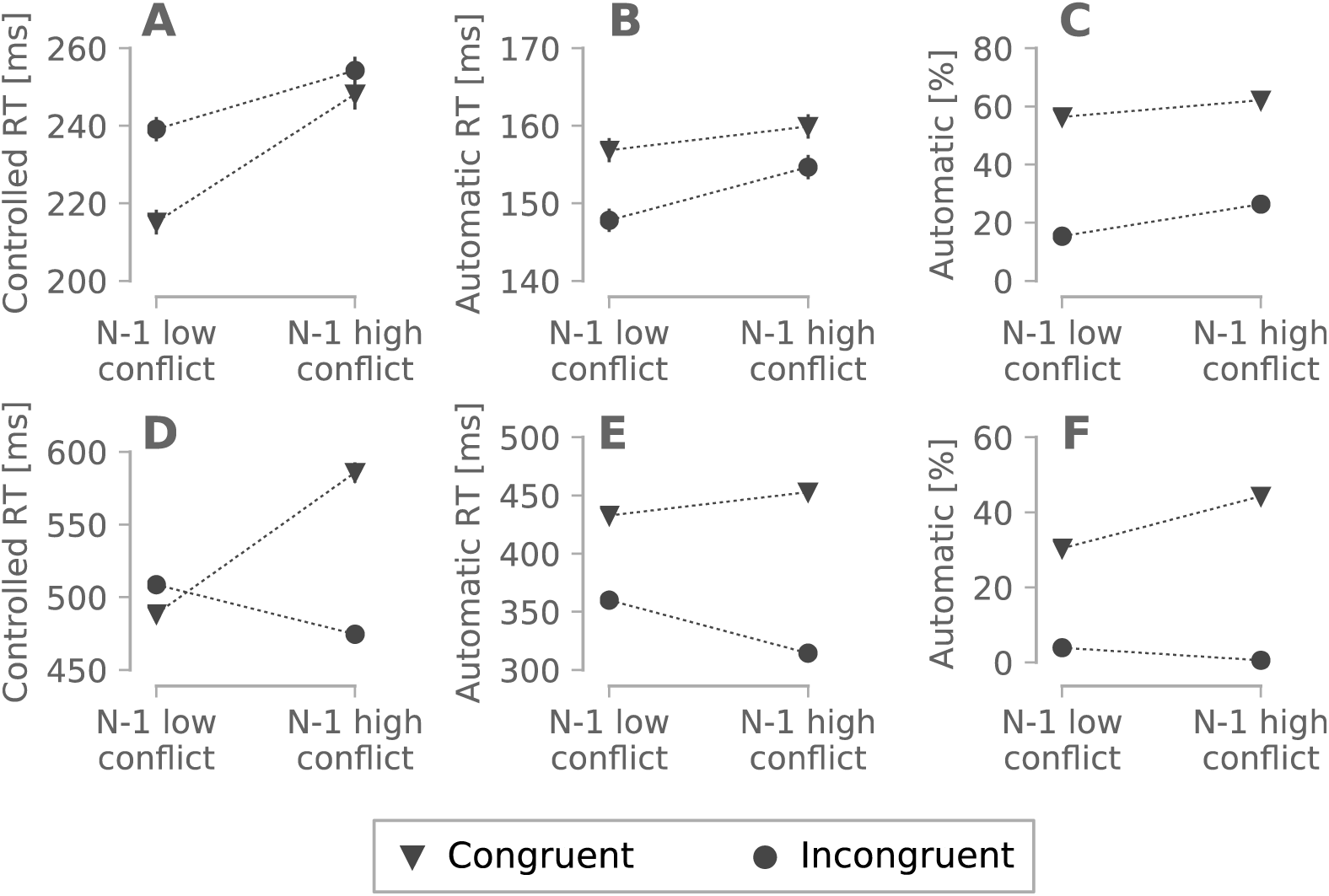
Model based analyses. Top row: Antisaccade task. Bottom row: Simon task. **A)** Reaction time (RT) of *correct* controlled congruent and incongruent responses in the antisaccade task. **B)** RT of automatic responses in congruent and incongruent trials in antisaccade task. **C)** Probability of an automatic response in the antisaccade task. **D-F)** Similar to A-C in the Simon task. Error bars display the standard error of the mean.

By contrast, congruent responses generated by the controlled process were much slower after a high conflict trial than after a low conflict trial (Δ = 97*ms*;*P* < 10^―5^; Fig. 4D). In other words, conflict-induced adaptation specifically slowed controlled congruent responses and this large inhibitory effect explained, to a large extent, the inversion of the Simon effect after high conflict trials. In addition, incongruent responses were facilitated after high conflict trials, as evidenced by reduced RT (Δ = 33*ms*, *P* < 10^―5^; Fig. 4D). Thus, conflict adaptation in the Simon task was expressed both as inhibition of congruent responses as well as facilitation of incongruent responses.

The antisaccade task displayed an analogous effect. Controlled congruent responses were considerably slower after high conflict trials, compared to low conflict trials (Δ = 32*ms*;*P* < 10^―5^; Fig. 4A). Importantly, this was not directly observable in the empirical mean RT because automatic responses were the dominant component in the congruent condition (61% of all responses; Fig. 4C), masking the contribution of controlled responses. However, in contrast to the Simon task, incongruent responses were also slower after the high conflict condition (Δ = 14*ms*; *P* < 10^―5^; Fig. 4A).

The previous analysis explains the congruency effect and conflict adaptation across tasks and conditions but it does not explain its *time course*, i.e., it does not explain the negative slopes in the delta plot in the Simon task. Because the bulk of conflict adaptation was caused by the inhibition of controlled congruent responses, we investigated them in isolation by removing automatic responses from the model predictions. In other words, we used SERIA to predict the distribution of controlled responses in the absence of automatic responses (see Methods). In addition, we plotted the time course of the congruency effect when only the distributions of *controlled* responses were taken into account.

Fig. 5 demonstrates that negative slopes in the delta plots are neither unique to the Simon task nor to trials following high conflict conditions. Instead, the variance of congruent controlled responses was higher than the variance of incongruent responses in all conditions and tasks, leading to negative delta plot *slopes* in this analysis (34). However, this effect was masked by the contribution of automatic responses, especially in the antisaccade task. Hence, conflict adaptation led to negative delta plots in post-conflict trials in the Simon task, but negative *slopes* were a more general effect caused by the high variance of congruent controlled responses.

**Figure 5:**
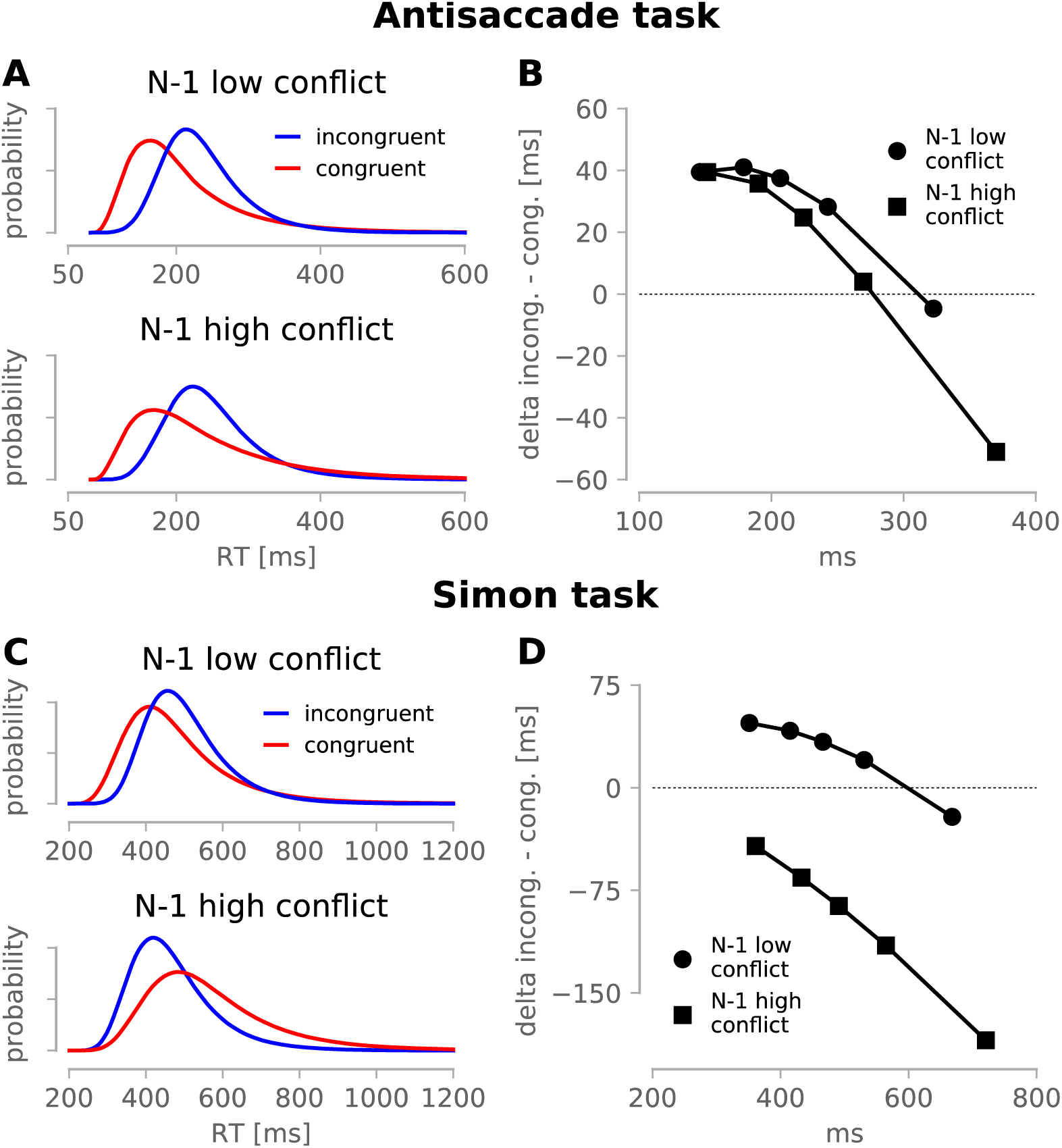
Model based analyses of controlled responses. Top row: Antisaccade task. Bottom row: Simon task. **A)** Normalized reaction time (RT) distribution of controlled responses in the antisaccade task. **B)** Delta plots between the distribution of controlled response in the antisaccade task. **C & D)** Similar to A and B, but for the Simon task. Congruent responses were more variable than incongruent responses in both tasks. This led to delta plots with negative slopes in all conditions. However, only after high conflict trials in the Simon task, the congruency effect was negative in all time bins.

However, the main explanation for conflict adaptation in the Simon task was the inhibition of congruent controlled responses across all time bins, which is evidenced by Fig. 5D as negative delta plots. In other words, conflict adaptation in the Simon task manifests as an overall inhibition of congruent controlled responses, which is separable from the negative *slope* of the delta plots. This is absent in the antisaccade task, where no significant conflict adaption was detected.

The success of SERIA in fitting the data from these two tasks begs the question whether these paradigms, both considered well validated response inhibition or interference control tasks, measure similar biological functions. If that were the case, we expected that the model-based estimates would correlate across tasks. However, neither the percentage of automatic responses nor their RT, as estimated by SERIA, were significantly correlated after correcting for multiple comparisons (Supp. Table 9). In a purely behavioral analysis, there was a significant but weak correlation between ER on incongruent trials (*r* = 0.208, *P* < 10^―3^ across tasks (Supp. Table 10), but this correlation should be interpreted with caution because subjects rarely made errors in the Simon task (mean ER < %5) and in 30% of cases did not make any at all (Supp. Fig. 2).

Three main conclusions can be drawn from Exp. 1. First, the same principles can be used to describe and predict behavior in the antisaccade and Simon tasks. Specifically, congruent responses can be generated by a fast but automatic process or by a controlled but slow process. Moreover, response inhibition arbitrates between these two components in a time dependent fashion.

The second main conclusion is that after high conflict trials, controlled congruent responses are inhibited as demonstrated by post-conflict slowing. In the Simon task, this led to a negative congruency effect. In the antisaccade task, SERIA also revealed a slowing of congruent responses. However, the delta plot analyses (Fig. 5 B & D) reveals the key difference between tasks: there was generalized slowing in the Simon task but not in antisaccade task, which explains the absence of post-conflict slowing in the latter paradigm. Complementary to the inhibition of congruent responses, incongruent responses were facilitated in the Simon task, but not in the antisaccade task.

Finally, our analysis suggests that one of the most remarkable properties of the Simon task — the negative time course of the congruency effect — is likely not specific to this task and not only caused by the inhibition of automatic responses (21–23). Rather, negative slopes are partly caused by the high variability of congruent controlled responses. However, this effect was masked in the antisaccade task, where most congruent responses were automatic. An analogous phenomenon occurred after low conflict trials in the Simon task, but this effect was again hidden by the contribution of automatic responses.

These findings set the stage for investigating the role of GABA-A signaling in response inhibition and conflict adaptation using SERIA. The former is reflected by the probability and latency of automatic responses. If response inhibition is facilitated by GABA-A, enhanced GABA-A activity should lead to *fewer* automatic responses with shorter latencies, as the inhibitory process would only fail to stop the fastest automatic actions. This hypothesis can be directly tested using computational modeling, as the effect of GABA-A on voluntary responses can be disentangled from the causes that may lead to decreased performance in the Simon and antisaccade task under benzodiazepines. Increased cognitive control should manifest as higher conflict adaptation expressed by either stronger inhibition of congruent controlled responses or by facilitation of incongruent responses after high conflict trials. In Exp. 2, we investigated the effect of lorazepam (a nonselective positive allosteric modulator of the GABA-A receptor) on the Simon and antisaccade tasks in a new sample.

### Experiment 2

Data from Exp. 2 were previously reported in (33). The same tasks were administered in Exp. 1 and 2 but in Exp. 2, each subject (N=50) participated in three sessions, in which either placebo or lorazepam (0.5 or 1.0mg) were administered. We excluded subjects from the final analysis if any of the three sessions would be excluded by the criteria used in Exp. 1. This left 38 valid subjects in the antisaccade task and 46 in the Simon task. As detailed in Supp. Fig. 1 & 3, we replicated all behavioral and modeling findings from Exp. 1 in this independent sample. Thus, we focus here only on the effects of lorazepam.

We first analyzed RT and ER using classical statistics, recapitulating the findings in (33). RT and ER increased with dose in both tasks (Table 1; see Fig. 6). In terms of RT, lorazepam did not have a significant effect on either the antisaccade cost or the Simon effect (no significant two- or three-way interaction between drug, trial type and conflict; Supp. Table 5 & 6). The effect of lorazepam on ER in the antisaccade and Simon tasks was modulated by trial type and conflict (P=0.019 and P=0.022 respectively; Supp. Table 7 & 8).

**Figure 6:**
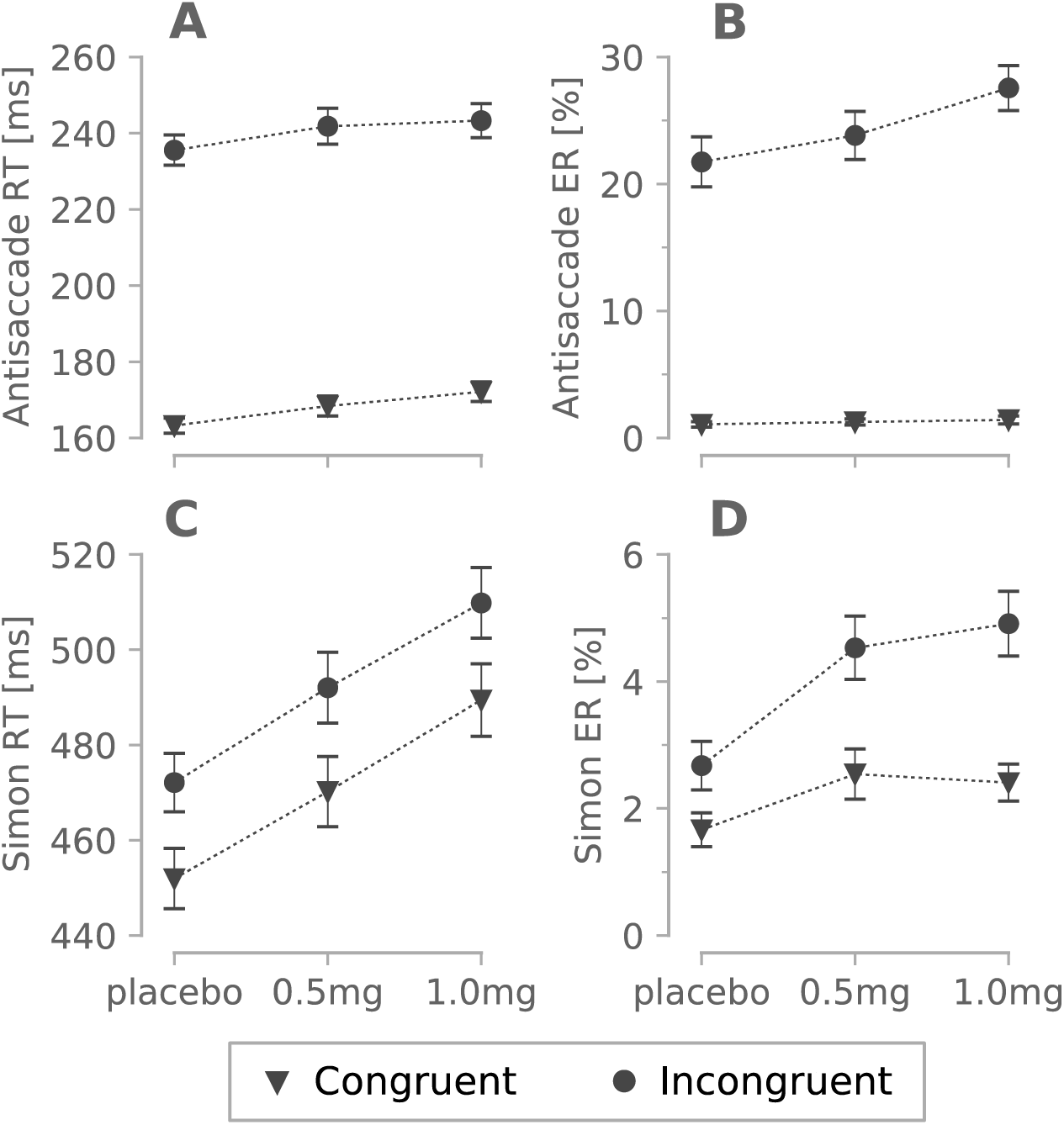
Effect of lorazepam (0.5 or 1.0mg) on reaction times (RT) and error rates (ER) in the antisaccade and Simon tasks. **A)** RT in the antisaccade task. **B)** ER in the antisaccade task. **C)** RT in the Simon task. **D)** ER in the Simon task. Error bars display the standard error of the mean.

**Table 1.** Lorazepam related behavioral effects. Error rate (ER) and mean reaction time (RT) increased in a dose dependent fashion.

|  | Antisaccade task |  |  |  | Simon task |  |  |
| --- | --- | --- | --- | --- | --- | --- | --- |
|  | Congruent trials |  |  |  |  |  |  |
|  | Placebo | 0.5mg | 1.0mg |  | Placebo | 0.5mg | 1.0mg |
| ER [%] | 1.0 | 1.2 | 1.4 |  | 1.6 | 2.5 | 2.4 |
| RT [ms] | 163 | 168 | 172 |  | 451 | 470 | 489 |
|  | Incongruent trials |  |  |  |  |  |  |
|  | Placebo | 0.5mg | 1.0mg | P | Placebo | 0.5mg | 1.0mg |
| ER [%] | 21.7 | 23.8 | 27.5 |  | 2.6 | 4.5 | 4.9 |
| RT [ms] | 235 | 241 | 243 |  | 472 | 492 | 509 |

To better understand the effect of lorazepam on response inhibition and conflict adaptation, we applied SERIA to trial-by-trial RT. Lorazepam impaired controlled responses as reflected by higher RT across tasks and trial types (see Fig. 7 & 8). Supp. Table 13 displays the detailed breakdown and statistical analysis of these effects. The latency of automatic responses increased in a dose dependent fashion in the antisaccade (*P* < 10^―3^) and Simon tasks (*P* = 0.11). Lorazepam only significantly raised the number of automatic responses at the 0.5mg dose in the Simon task (*P* < 10^―3^). Thus, rather than enhancing response inhibition, lorazepam impaired it in terms of the RT and the probability of automatic responses in the Simon task.

**Figure 7:**
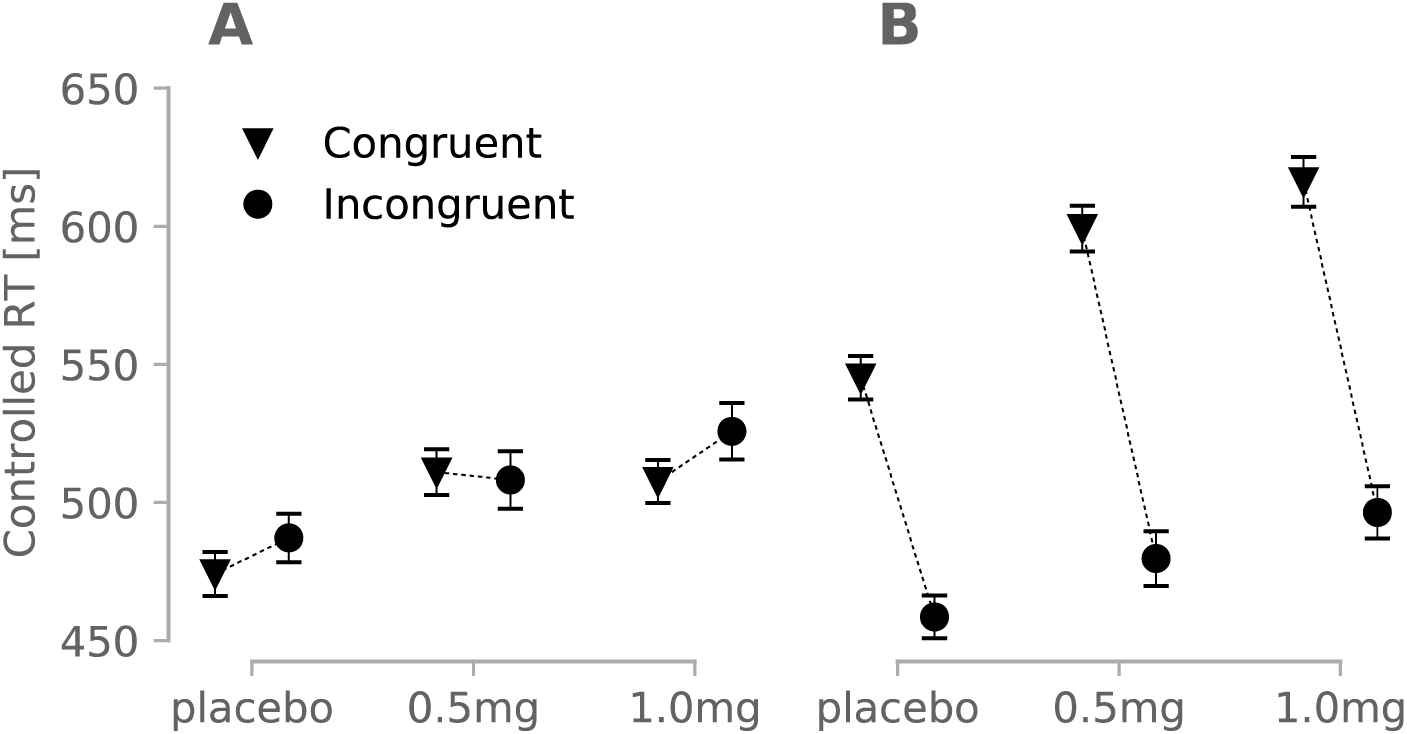
Mean RT of controlled responses in the Simon task. Reaction time (RT) increases as a function of dose. **A)** Trials following the low conflict (congruent) condition. **B)** Trials following the high conflict (incongruent) condition. Response interference was boosted by lorazepam, measured as the difference between controlled congruent and incongruent responses after high conflict trials.

**Figure 8:**
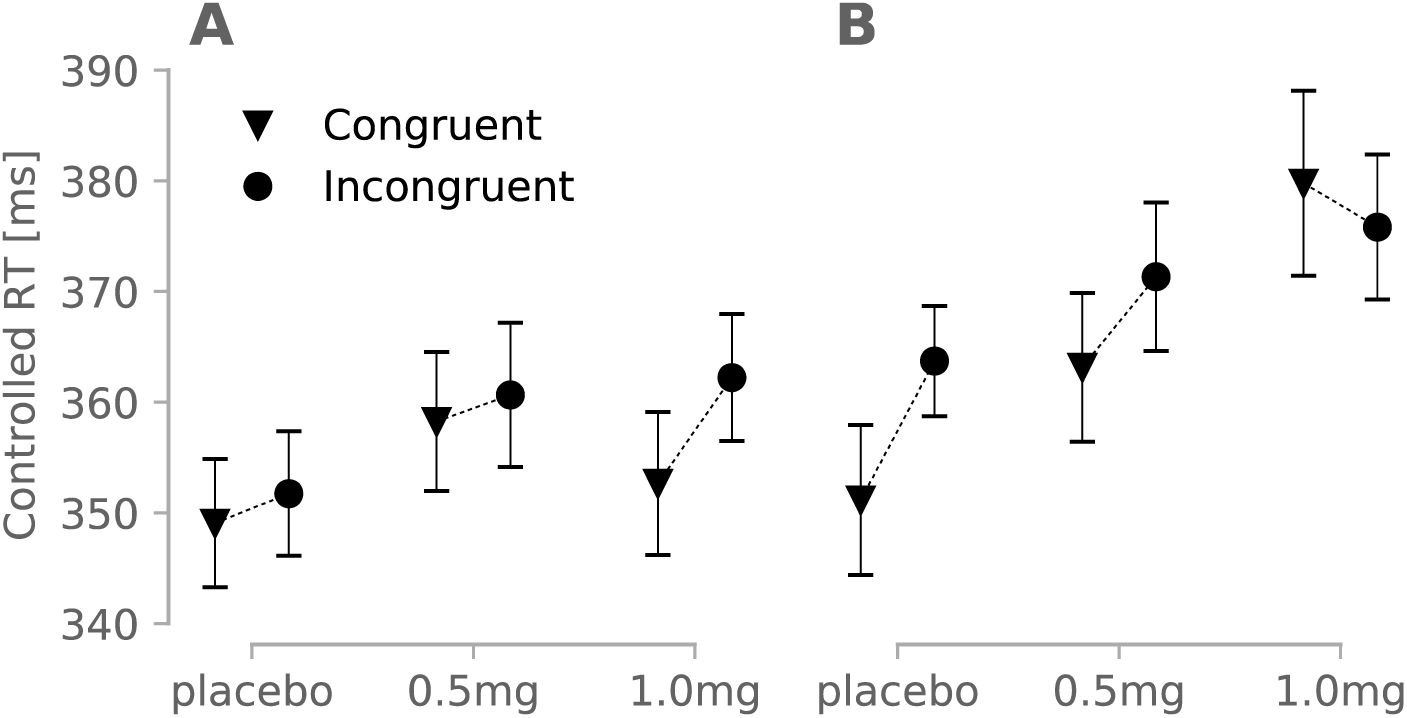
Mean reaction time (RT) of controlled responses in the antisaccade task. A) Trials following the low conflict (congruent) condition. B) Trials following the high conflict (incongruent) condition. Lorazepam increased congruent and incongruent RT (*P* < 10^―5^).

One of the main findings from Exp. 1 is that the Simon effect turned negative after high conflict trials because congruent controlled responses were inhibited following high conflict trials. Can heightened GABAergic signaling enhance conflict adaptation by improving the inhibition of congruent controlled responses? Indeed, lorazepam led to a dose dependent slowing of controlled congruent responses after high conflict trials (see Fig. 7B) as evidenced by a three-way interaction between the factors drug, previous trial conflict level and trial type (*P* = 0.01). To understand this three-way interaction, we split congruent and incongruent trials and tested for interactions between drug and conflict. Incongruent controlled responses were facilitated after high conflict trials as compared to those following low conflict trials (Δ = ―29*ms*;*P* < 10^―5^), but there was no significant interaction between lorazepam and conflict on those trials (*P* = 0.992). By contrast, congruent responses were slower after high conflict trials compared to low conflict trials (Δ = 89*ms*;*P* < 10^―5^), and this effect was modulated by lorazepam (*P* = 0.001). Thus, lorazepam boosted inhibitory conflict adaptation in congruent trials, but did not have a significant impact on incongruent controlled responses.

In the antisaccade task (Fig. 8), lorazepam increased the mean RT of controlled responses (*P* < 0.001), but there was no significant three-way interaction between the factors drug, trial type and previous trial conflict (*P* = 0.176). When the effect of conflict level in the previous trial was analyzed separately on congruent and incongruent trials, lorazepam did not significantly interact with conflict level on incongruent (*P* = 0.901) but it did so on congruent trials (*P* = 0.046). Although not strong, the latter effect pointed in the same direction as in the Simon task. As suggested by Fig. 8B, this effect was particularly salient at the highest dose (1mg lorazepam).

## Discussion

The concept of *response inhibition* is a pivotal construct in cognitive neuroscience and psychology. A formal model of this construct should be compatible with two broad observations: (i) some actions can be triggered in an automatic, stimulus driven manner and (ii) a more deliberative, yet slower decision process can generate a larger set of behaviors. The automatic process is tuned to situations that are either stereotypical or that require fast responses. By contrast, the second decision process allows for a richer behavioral repertoire in accordance with goals, feedback, and changes in the environment. While these observations are far from contentious, it is still an open question how these processes interact with each other.

SERIA formalizes the notion that response inhibition is the mechanism that mediates between controlled and automatic responses, and that, in two-alternative force-choice decision tasks, response inhibition is a time dependent process. Our model can be derived from these general premises by adding the assumption that the decision process between controlled actions is also a race to threshold. This is a generalization of the traditional and successful horse-race model (27) of response inhibition used in the context of the stop signal task. Because of its simplicity, SERIA can be formulated in an analytical manner and fitted to trial-by-trial RT and actions. This is in contrast to the common approach of fitting histograms or cumulative density functions (35–37).

Surprisingly, the interplay between conflict adaptation and response inhibition has not been explored in detail so far, even though both functions are seen as components of executive control and have been studied with similar paradigms. As suggested before, one reason for this is the lack of analytical computational models that bring together both functions under a single roof. One of the main results of this study is to show that a single computational model (SERIA) can explain the interplay of response inhibition and conflict adaptation in individual subjects. Specifically, our two data sets demonstrated that SERIA can fully account for RT and ER in the Simon and antisaccade tasks. Indeed, not only was SERIA able to accurately predict RT distributions (Fig. 3A&D), mean RT and ER (Supp. Fig. 1), but it also captured the time course of the congruency effect as visualized in the delta plot analysis (Fig. 3B,C,E,F).

SERIA identified how heightened cognitive control and conflict adaptation manifest after a high conflict trial: *controlled congruent responses* are slower after incongruent trials, both in the Simon and antisaccade tasks. This effect was not obvious in the antisaccade task because the large number of automatic responses masked changes in controlled congruent responses. However, our in-depth model-based analysis unveiled this change. In addition, high conflict trials facilitated incongruent responses in the Simon task, while the opposite effect was observed in the antisaccade task.

The second experiment allowed us to investigate whether response inhibition and conflict adaptation are mediated by GABA-A signaling using the benzodiazepine lorazepam. Regarding response inhibition, lorazepam increased the latency of fast responses. However, this effect was not specific as it was shared by controlled responses and was to be expected from the sedative effect of benzodiazepines (38, 39). Interestingly, we have previously shown (40) that dopamine does not strongly mediate response inhibition in the antisaccade task, whereas cholinergic modulation leads to a dose dependent increase in the number of inhibition failures.

Lorazepam had a dose dependent effect on conflict adaptation as shown by the slowing of congruent controlled responses after high conflict trials in the Simon task. In other words, GABAergic signaling boosted cognitive control in the Simon task.

Previous studies in other paradigms have suggested that cognitive control manifests as amplification of task relevant information and not inhibition of task irrelevant stimuli (41). While our results are in contrast to this study, it is now clear (6) that findings related to cognitive control are largely domain-specific and do not easily generalize across paradigms. For example, in our study, incongruent responses were facilitated after high conflict trials in the Simon task, but not in the antisaccade task. Nevertheless, here we offer evidence that cognitive control can be enhanced by conflict-induced GABAergic inhibition.

Our findings invite the question whether conflict adaptation is impaired in generalized anxiety, for which benzodiazepines are a common second-line treatment. Indeed, there is strong evidence that generalized anxiety has a detrimental effect on cognitive control (reviewed in (42)). More specifically, conflict adaptation but not conflict monitoring appears to be blunted in subjects with high generalized anxiety. For example, (43, 44) found impaired conflict adaptation using the emotional Stroop task (45). It is worth noting that more recent studies (46, 47) have found physiological differences in conflict adaptation between controls and generalized anxiety individuals, but behavioral evidence has not always been positive. However, these studies did not control for medication. Thus, one might hypothesize that benzodiazepines have a positive effect on conflict adaptation thereby normalizing this function in generalized anxiety. Interestingly, anxiety induced by the neuropeptide cholecystokinin-tetrapeptide (CCK4) leads to overactivation of the anterior cingulate cortex, an area critical for conflict adaptation (4, 5), and this effect can be prevented by 1mg of the benzodiazepine alprazolam (48). To our knowledge, direct evidence of the role of benzodiazepines in conflict adaptation in anxiety is still lacking.

### Other models

SERIA can be seen as a formal version of the activation-suppression model (22, 23). It offers a unified conceptual and formal account of response inhibition in the antisaccade and Simon tasks and is compatible with the horse race model of the stop-signal task. However, SERIA and the activation suppression model offer different explanations for the negative slope of the delta plots in the Simon task. The activation suppression model asserts that negative slopes are caused by the time dependent inhibition of congruent responses, which is weak at short latencies but strengthens over time.

Our computational analysis offers a subtler explanation. According to SERIA, in the antisaccade and Simon tasks, congruent controlled responses are more variable than incongruent (controlled) responses. This leads to negative slopes in the delta plot analysis when the contribution of automatic congruent responses is removed from the distribution of congruent responses (see Fig. 5). The weight of this effect is modulated by the ratio of *controlled* and *automatic* congruent responses, as the latter have shorter latencies and are less variable than controlled responses. In the antisaccade task, automatic responses are dominant, leading to delta plots with positive slopes independently of the conflict level in the previous trial. In the Simon task, automatic responses are less common, leading to directly observable negative slopes after high conflict trials.

In summary, three factors together explain the negative slopes in the Simon task: the larger variability and latency of controlled congruent responses (a factor also present in the antisaccade task), the relatively low number of automatic responses, and conflict adaptation, expressed as faster incongruent responses after high conflict trials.

Other computational models have been proposed for the antisaccade task (discussed in detail in (26,49,50) and the Simon task (24, 25). Historically, the main constraint on computational accounts of the Simon effect has been the ability to simulate its negative slope as a function of time because this was initially understood as evidence that controlled and automatic processes were active in the Simon task. While this notion was soon abandoned (34), as radically different models can simulate negative slopes in delta plots (25), this constraint still plays an important role.

Among the models that have been previously proposed, a particularly interesting approach is the extension of the drift-diffusion model (DDM) suggested by (37), which simulates automatic responses through an additive ‘bump’ in a linear diffusion process. Through this extension, Ulrich and colleagues could simulate the time course of the Simon effect without introducing a third inhibitory process, at the cost of losing analytical tractability and the ability to fit subject-by-subject responses. Currently, it is not possible to formally compare this extension of the DDM with SERIA, as only SERIA has a tractable generative form. Nevertheless, it remains an open question whether an independent stopping process is needed to account for response inhibition.

### Is response inhibition a unitary construct?

Despite of the structural similarities between the two tasks captured by SERIA, there were no significant correlations between parameter estimates across them. This negative finding reflects the accumulating evidence that the psychological construct of ‘response inhibition’ is heterogeneous and does not encompass a single executive function (51–53). Rather, the success of SERIA in capturing behavior under the two distinct tasks implies convergent mechanisms that depend on different biological functions.

## Conclusion

This study provides a novel and comprehensive account of the congruency effect in the Simon and antisaccade tasks, its time course, and how this effect interacts with conflict adaptation. Our account is supported by formal model comparison and highly accurate model fits, e.g., of the congruency effect, generated from trial-by-trial fits. We provide evidence that in the Simon task, conflict adaptation manifests both as facilitation of incongruent responses and inhibition of congruent responses after high conflict trial. Importantly, for the first time, we show that conflict induced response inhibition and conflict adaptation are modulated by GABA-A signaling. Our finding that lorazepam differentially impacts on response inhibition and conflict adaptation suggests that positive allosteric modulators of the GABA-A receptor do not affect cognitive control equally but modulate its component processes in different ways: conflict adaptation is facilitated by enhanced GABAergic signaling, whereas response inhibition is impaired in an unspecific manner.

## Methods

### Experimental procedure

The experimental procedure and data from Exp. 2 have been reported previously in detail (33). Thus, we briefly summarize the protocols emphasizing the difference between Exp. 1 and 2. All experimental procedures were approved by the research ethics committees of the Department of Psychology (Exp. 1) and the Faculty of Medicine (Exp. 2) at the University of Bonn and followed the Declaration of Helsinki.

### Experiment 1

In Exp. 1, 164 healthy subjects (mean 23 ± 3 years of age; 81 females) performed the antisaccade and Simon tasks. The order in which the tasks were administered was pseudorandomized across subjects. Each task consisted of 100 congruent and incongruent trials displayed in random order. Data from Exp. 1 are first reported here.

Trials in the antisaccade task started with a central fixation stimulus (random duration of 1000-2000ms) followed by a peripheral cue displayed for 1000ms either on the left or right side of the screen (±10.3°). Subjects were instructed to saccade either to the cue or in the opposite direction, depending on the color of the central fixation (blue or yellow). Eye gaze was measured with an EyeLink 1000 (SR Research, Canada) at 1000 Hz sampling rate. RT was defined as the latency between the presentation of the peripheral cue and the first saccade following this. Saccades with latency lower than 80ms were considered invalid.

In the Simon task, subjects were instructed to press a left (‘x’) or right key (‘,’) on a QWERTZ keyboard depending on the color of a circular cue presented for 1500ms on the left or right side of the screen following a 500ms central fixation cross. RT was defined as the latency between cue presentation and the first key press. In both tasks, error rate (ER) was calculated as the ratio between number of valid incorrect responses and total number of valid responses.

### Experiment 2

Exp. 2 (described in (33) consisted of the same tasks as in Exp. 1. A new sample of N = 50 healthy volunteers took part (mean 24 ± 3 years of age; 27 females). However, subjects were administered placebo and lorazepam (0.5 or 1.0 mg) across three different sessions in a within-subject, double-blind, randomized design. The order in which the tasks were administered was randomized across subjects but kept constant across sessions.

### Data processing

Details of data preprocessing can be found elsewhere (33). Subjects with fewer than 65% of valid trials or more than 80% ER in any session were excluded from the final analysis.

### Modeling

Responses (congruent/incongruent) and their respective RT were modeled with SERIA (26) which posits that actions are the outcome of the competition between four race-to-threshold processes. First, automatic responses are generated by a fast process, which we call the automatic process. In the antisaccade task, this process can only generate saccades toward the cue (prosaccades). In the Simon task, automatic responses are always congruent responses toward the location of the cue (e.g., right button presses for right stimuli, irrespective of the color of the cue). Automatic responses can be stopped by a latent, unobservable inhibitory process, when the latter hits threshold before the automatic process. When this happens, the second race between congruent and incongruent *controlled* responses determines which response is generated. This second decision is modeled with two race-to-threshold processes that compete against each other. The mathematical details of the model can be found in (26). Each of these processes is parametrized by its mean threshold hit time and the corresponding variance.

In addition to these 8 parameters (2 per process), SERIA uses three auxiliary parameters: the no-decision time which reflects neuronal transmission delays until the race of the first unit starts, the probability of low latency outliers, i.e., of reactions whose RT is below the no-decision time, and an extra delay associated with the start of the race between controlled responses. Thus, in its more general version, SERIA has 11 parameters.

The antisaccade and Simon tasks were modeled independently. In each task, trials from each session were divided into congruent and incongruent trials. These were split up into those that followed congruent (low-conflict) and incongruent (high-conflict) trials. Hence, each session consisted of 4 different conditions, each of which could be modeled with a potentially different set of parameters, i.e., with 44 different parameters.

To constrain the model and to avoid overfitting, the auxiliary parameters were assumed to be constant across the 4 conditions. In addition, the parameters of the automatic process were kept constant across the four conditions. Otherwise, these parameters would be strongly correlated with the parameters of the inhibitory process. Moreover, the parameters of the *incorrect controlled response* were the same across all conditions. In other words, the distribution of the hit times of the incorrect controlled response (e.g., a controlled congruent response on an incongruent trial) was kept constant irrespective of trial type and the conflict level of the previous trial. Finally, the parameters of the inhibitory unit on congruent switch and repeat trials were fixed to the same values. This yielded a total of 20 free parameters, i.e., degrees of freedom, per experimental block comprising the four conditions.

We evaluated the possibility that this model overfits the data by comparing it to a simpler model in which congruent responses are always automatic, fast responses but with the same number of parameters. Thus, we could verify if a structurally simpler model, with a comparable number of parameters could parsimoniously explain subjects’ performance. As shown in Supp. Table 1 & 2, SERIA was more parsimonious and accurate than a reference model with equal number of parameters but less structural flexibility.

In Exp. 1, SERIA was fitted using the Metropolis-Hastings algorithm in combination with a hierarchical model. This was used to estimate the prior mean and variance of subject-specific parameters based on the population distribution (i.e., an empirical Bayesian procedure). For Exp. 2, this model was enriched by modeling the effect of lorazepam, i.e., in addition to the population mean and variance, we accounted for the effect of the two doses. Moreover, we modeled subject specific intercepts as random effects. This model provides empirically motivated priors, and we have used it in a variety of previous studies; for details, see (30, 40).

The convergence of the Metropolis-Hastings algorithm was evaluated with the *R* statistic (54), such that at most 2% of all the model parameters were allowed to cross the 1.1 threshold (*R* > 1.1), commonly used to assert convergence.

Fits and delta plots were generated from the posterior predictive distribution of each subject. In other words, we used the parameter estimates to approximate the conditional density

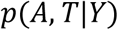

where *A* ∈ {*congruent,incongruent*} is the action generated in a trial, *T* ∈]0,∞[its RT and *Y* represents the empirical data from a subject.

In practice, the posterior predictive distribution can be estimated by averaging out the likelihood computed from samples collected using MCMC, as

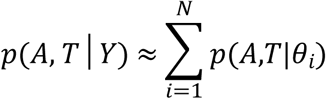

where *θ*_*i*_ are samples from the posterior *p*(*θ*|*Y*).

To analyze the distribution of voluntary congruent and incongruent responses in isolation from congruent responses (see Fig. 5), we use the posterior parameter estimates to compute the distributions:

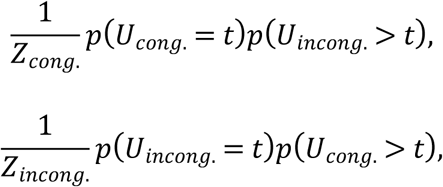

where

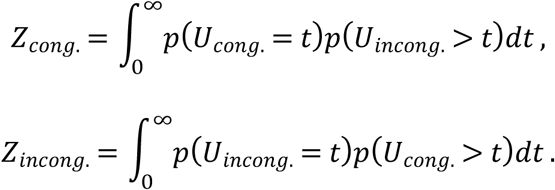

when *U_cong_*. and *U_incong_*. are the hit time of the controlled congruent and incongruent decision processes. These distributions predict the RT of congruent and incongruent responses in case no automatic response would have taken place.

We also report mean hit time of the controlled congruent and incongruent responses defined as

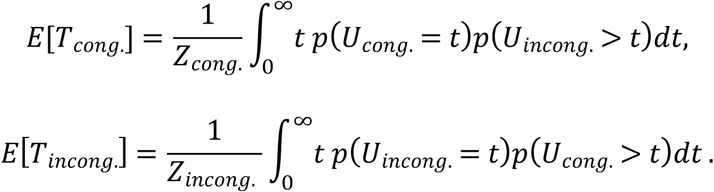

The probability of an automatic response (which we have called inhibition failure in previous studies) is defined as the probability that the automatic process hits threshold before any other unit.

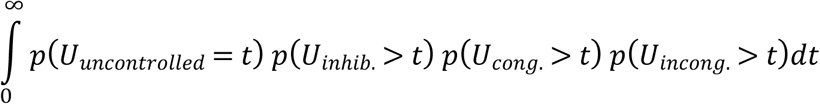

All methods described here are available in the open source TAPAS toolbox. The code used for the analysis is openly available at www.translationalneuromodeling.org/tapas.

### Statistical analysis

Behavioral data as well as model parameters were analyzed using generalized linear mixed-effect models (GLME), implemented in the R (3.6.1) statistical package. The independent variables were conflict in the previous trial with levels high (incongruent) and low (congruent), trial type (levels congruent and incongruent) and subject, treated as a random effect. In Exp. 2, we also modeled the effect of lorazepam with levels placebo, 0.5mg, 1.0mg treated as a categorial regressor. For ER, a logistic regression model was used, and significance was assessed with Wald tests. Probabilities estimated using SERIA were analyzed using Beta regression models implemented in the package glmmADMB and significance was again assessed using Wald tests. For RT, F-tests were used in combination with the Satterthwaite approximation to the degrees of freedom.

## Acknowledgements

We thank Klaas E. Stephan for his comments on the manuscript.

## Supporting Information

**Supplementary Figure 1:**
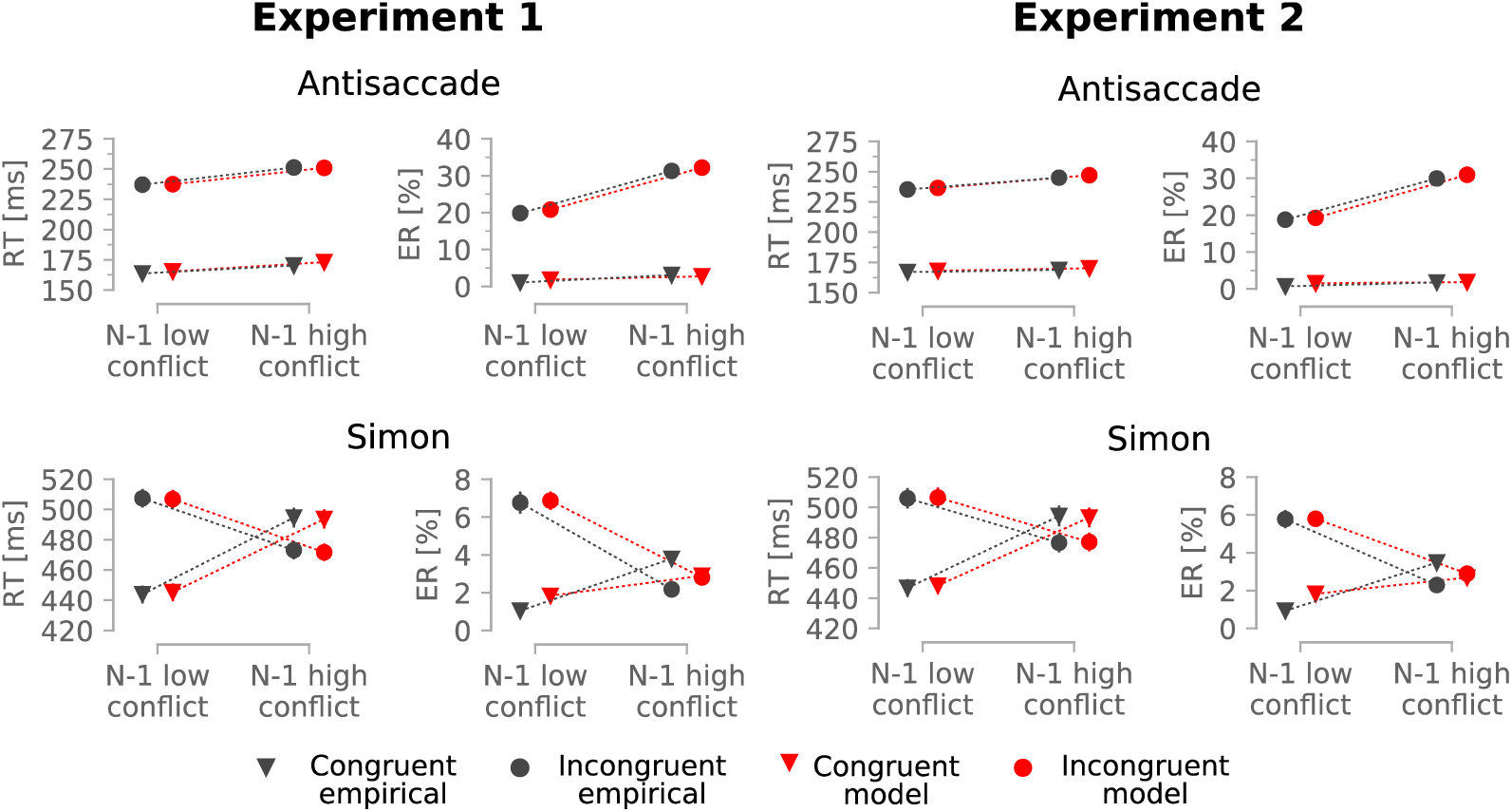
Empirical and predicted reaction time (RT) and error rate (ER) in Exp. 1 and 2. All effects from Exp. 1 were replicated in Exp. 2. The model accurately reproduced the empirical RT and ER. Error bars display the standard error of the mean.

### Supplementary material 1 – Model comparison

The key assumption of the SERIA model is that congruent responses can be generated by either a fast, automatic process, or a slow but flexible component. We have shown previously (Aponte et al., 2017) that in some versions of the antisaccade task, RT are bimodally distributed and this constitutes qualitative evidence for the SERIA model. In the versions of the Simon and antisaccade tasks administered here, RT were unimodally distributed, which begs the question whether it is necessary to postulate two decision processes to explain RT and ER on congruent trials.

To answer this question, we compared SERIA to models in which all congruent responses are generated by a single process. Mathematical details for model comparison of different SERIA models can be found in (Aponte al., 2018). We report two metrics: the accuracy (or expected log likelihood), and the Watanabe Akaike information criterion (WAIC) (55), which is equal to the pointwise log likelihood (also called pointwise log predictive density) minus a penalization term that represents complexity. Two variants of each model were tested (see Supp. Table 1-2). In model *m*_1_ and *m*_2_, all congruent responses were generated by an uncontrolled process. In model *m*_1_, we allowed the parameters of controlled responses to be different depending on the four possible conditions (trial type x N-1 conflict level). In model *m*_2_, the parameters of the controlled responses were always equal across conditions. The difference between model *m*_3_ and *m*_4_ is that the variance of the inhibitory unit was kept constant across all conditions, whereas in model *m*_3_ it was different across congruent and incongruent trials. We introduced this model to further simplify model *m*_3_ and improve the convergence of the inference algortihm. All results reported here were obtained with model *m*_4_.

In general, SERIA fitted the data from both tasks more accurately and had higher WAIC than single process models with a comparable number of parameters. Thus, the structural flexibility inherent to dual process models explained RT distributions and ER better than single process models.

**Supplementary Table 1.** Model comparison in the antisaccade task Experiment 1. Four different models were evaluated based on their accuracy (expected log likelihood) and the Watanabe Akaike Information Criterion (WAIC). In model 1 and 2, congruent responses were generated by a single process. In model 2, (automatic) congruent responses were assumed to be identical in all conditions. This model was introduced because in models 3 and 4, all automatic responses (which are always congruent) were identical across conditions. In addition, in model *m*_4_ the variance of the inhibitory unit was not modulated by the conflict level on the previous trial. The highest WAIC is highlighted in bold.

|  | Model | #<br>parameters | Accuracy | WAIC |
| --- | --- | --- | --- | --- |
| $m_1$ | Single process | 27 | -19750 | -20655 |
| $m_2$ | Single process | 21 | -20579 | -21200 |
| $m_3$ | Dual process (SERIA) | 21 | -19117 | -19904 |
| $m_4$ | Dual process (SERIA) | 20 | -19086 | **-19871** |

**Supplementary Table 2.** Model comparison in the Simon task Experiment 1. Similar to Supp. Table 1. Bold entry indicates winning models, with almost identical WAIC.

|  | Model | #<br>parameters | Accuracy | WAIC |
| --- | --- | --- | --- | --- |
| $m_1$ | Single process | 27 | -43083 | -43783 |
| $m_2$ | Single process | 21 | -43940 | -44442 |
| $m_3$ | Dual process (SERIA) | 21 | -42754 | **-43379** |
| $m_4$ | Dual process (SERIA) | 20 | -42769 | -43384 |

In the Simon task, the two versions of SERIA used were very similar in terms of their accuracy and WAIC, relative to the size of the data set. This and all other results were replicated in Exp. 2 (Supp. Table 3 & 4). All analyses in the manuscript were based on model *m*_4_ as the differences in WAIC were small relative to the set size and it was the model with the fewest parameters). The key pharmacological result in Exp. 2 was significant regardless of the SERIA model used to estimate parameters.

**Supplementary Table 3.**
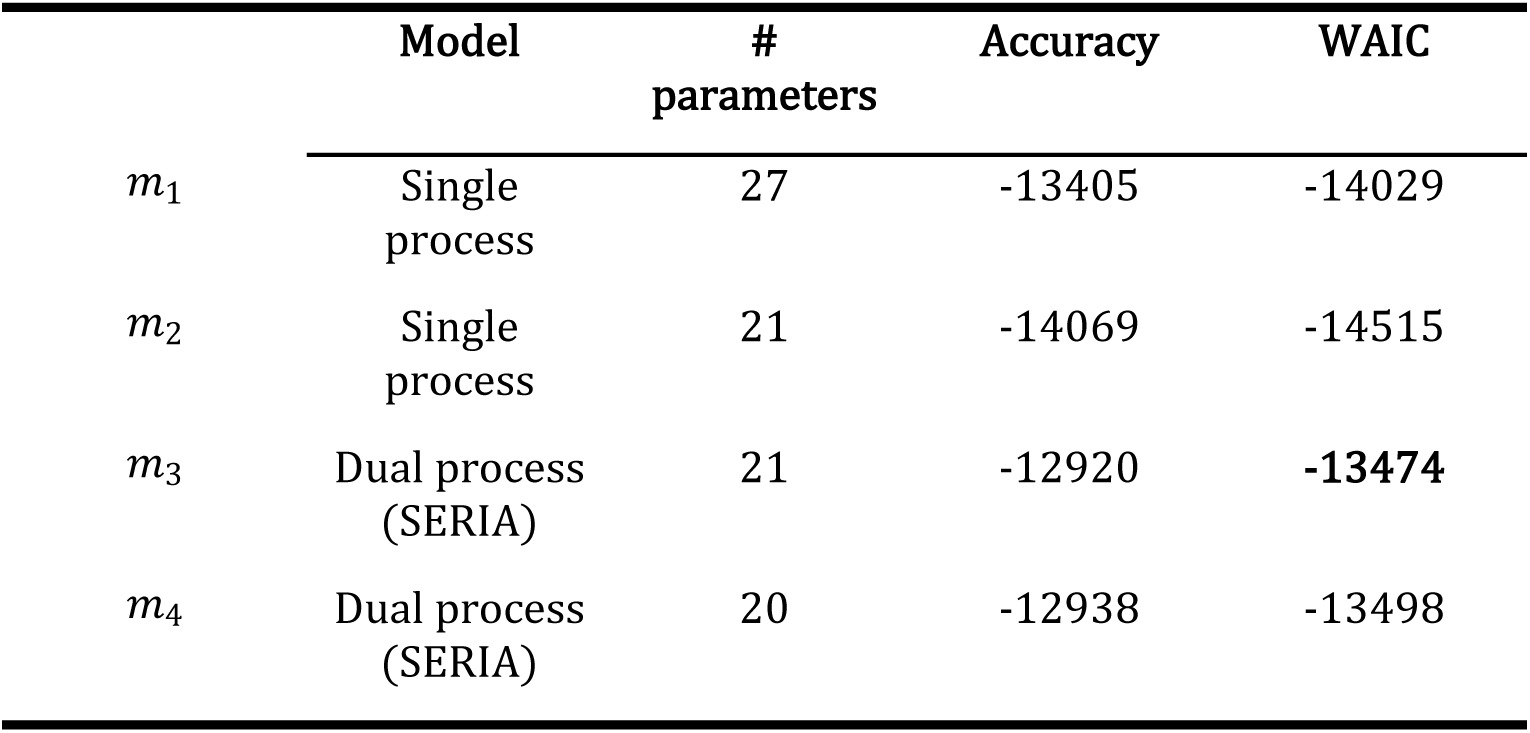
Model comparison in the antisaccade task Experiment 2. Similar to Supp. Table 1. Bold entry indicates the highest WAIC.

|  | Model | #<br>parameters | Accuracy | WAIC |
| --- | --- | --- | --- | --- |
| $m_1$ | Single process | 27 | -13405 | -14029 |
| $m_2$ | Single process | 21 | -14069 | -14515 |
| $m_3$ | Dual process (SERIA) | 21 | -12920 | **-13474** |
| $m_4$ | Dual process (SERIA) | 20 | -12938 | -13498 |

**Supplementary Table 4.**
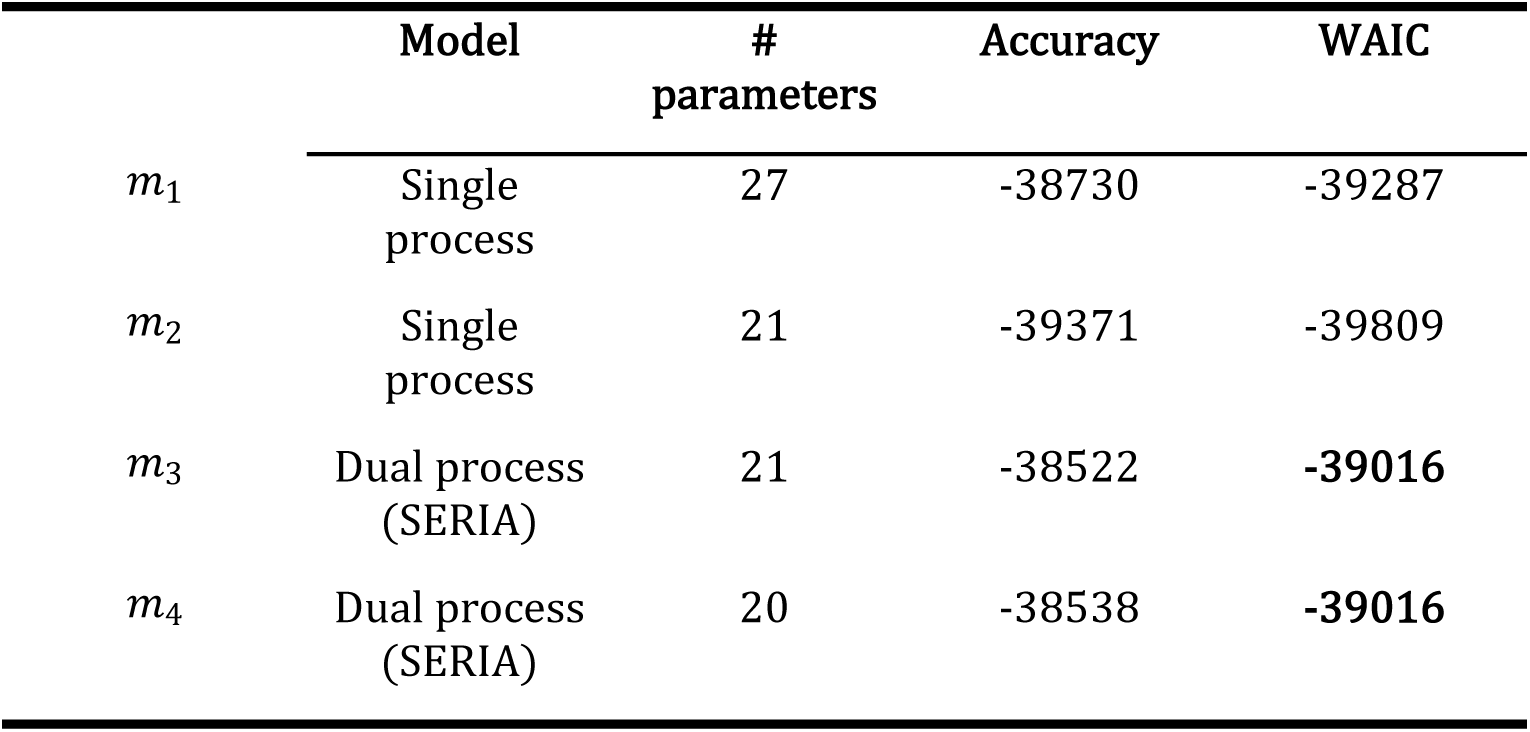
Model comparison in the Simon task Experiment 2. Similar to Supp. Table 1. Bold entry indicates winning models.

### Supplementary 2 - Experiment 2

Statistical analyses of mean RT and ER in Exp. 2 are displayed in Supp. Table 5-8 and summarized below. We do not report standardized effect sizes because these are not well defined for mixed effect models.

**Supplementary Table 5.** Antisaccade task Experiment 2 – Mixed effects ANOVA of the mean reaction time.

|  | NumDF | DenDF | F | P |
| --- | --- | --- | --- | --- |
| Trial type | 1 | 407 | 1398.216 | < 2.2e-16 |
| Conflict (N-1) | 1 | 407 | 8.917 | 0.003 |
| Dose | 2 | 407 | 6.457 | 0.002 |
| Trial type *<br>Conflict (N-1) | 1 | 407 | 4.211 | 0.041 |
| Trial type *<br>Dose | 2 | 407 | 0.114 | 0.893 |
| Conflict (N-1) *<br>Dose | 2 | 407 | 0.041 | 0.960 |
| Trial type *<br>Conflict (N-1) *<br>Dose | 2 | 407 | 0.103 | 0.902 |

**Supplementary Table 6.** Simon task Experiment 2 – Mixed effects ANOVA of the mean reaction time.

|  | Num DF | Den DF | F | P |
| --- | --- | --- | --- | --- |
| Trial type | 1 | 495 | 72.994 | <10<sup>-5</sup> |
| Conflict (N-1) | 1 | 495 | 13.884 | <10<sup>-5</sup> |
| Dose | 2 | 495 | 79.528 | <10<sup>-5</sup> |
| Trial type *<br>Conflict (N-1) | 1 | 495 | 250.079 | <10<sup>-5</sup> |
| Trial type *<br>Dose | 2 | 495 | 0.046 | 0.955 |
| Conflict (N-1) *<br>Dose | 2 | 495 | 1.040 | 0.354 |
| Trial type *<br>Conflict (N-1) *<br>Dose | 2 | 495 | 2.534 | 0.080 |

**Supplementary Table 7.**
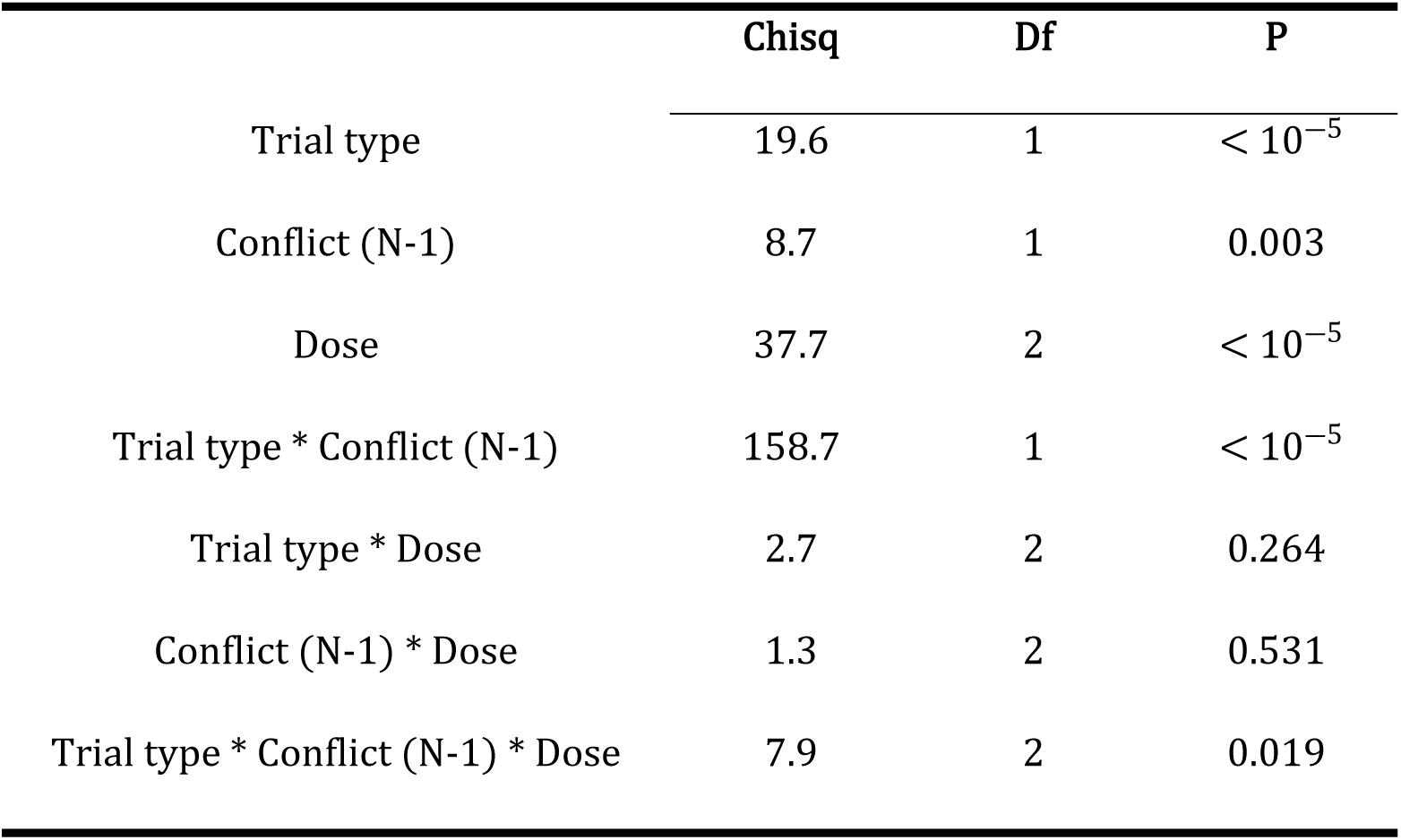
Antisaccade task Experiment 2 – Mixed effects ANOVA of the error rate.

|  | Chisq | Df | P |
| --- | --- | --- | --- |
| Trial type | 19.6 | 1 | $< 10^{-5}$ |
| Conflict (N-1) | 8.7 | 1 | 0.003 |
| Dose | 37.7 | 2 | $< 10^{-5}$ |
| Trial type * Conflict (N-1) | 158.7 | 1 | $< 10^{-5}$ |
| Trial type * Dose | 2.7 | 2 | 0.264 |
| Conflict (N-1) * Dose | 1.3 | 2 | 0.531 |
| Trial type * Conflict (N-1) * Dose | 7.9 | 2 | 0.019 |

**Supplementary Table 8.** Simon task Experiment 2 – Mixed effects ANOVA of the error rate.

|  | Chisq | Df | P |
| --- | --- | --- | --- |
| Trial type | 1298.4 | 1 | $< 10^{-5}$ |
| Conflict (N-1) | 202.9 | 1 | $< 10^{-5}$ |
| Dose | 40.7 | 2 | $< 10^{-5}$ |
| Trial type * Conflict (N-1) | 0.6 | 1 | 0.424 |
| Trial type * Dose | 0.3 | 2 | 0.855 |
| Conflict (N-1) * Dose | 4.2 | 2 | 0.120 |
| Trial type * Conflict (N-1) * Dose | 7.7 | 2 | 0.022 |

#### Behavioral analysis – Antisaccade task

Antisaccade mean RT (241ms) was higher than prosaccade RT (169ms; *P* < 10^―5^). The congruency effect (antisaccade cost) was higher after high conflict trials (76ms) compared to trials following low conflict trials (68ms). This yielded a significant interaction between the factor trial type and conflict (*P* = 0.042).

Regarding the mean ER, similar effects were observed as with RT. Antisaccade trials showed higher ER (6%) than prosaccade trials (1%; *p* < 10^―5^). Again, the congruency effect was higher after high conflict trials (27%) compared to low conflict trials (17%) but this was not reflected in a significant interaction between the factors trial type and conflict (*P* = 0.681).

#### Behavioral analysis – Simon task

In the Simon task, incongruent trials were on average 20ms slower than congruent trials (*P* < 10^―5^). In addition, trials that followed the high conflict condition were 10ms faster than trials following the low conflict condition (*P* < 10^―3^). Importantly, the Simon effect was significantly reduced after high conflict trial (―18*ms*), compared to low conflict trials (59*ms*) which yielded a significant interaction between previous trial conflict level and trial type (*P* < 10^―5^). We come back to this point in the next section.

The ER on incongruent trials was 1.8% higher than on congruent trials (*P* < 10^―5^). The congruency effect on trials following the high conflict condition (- 1%) was lower compared to trials that followed the low conflict condition (4%) and this interaction was significant (*P* < 10^―5^).

### Supplementary 3 – Correlations across tasks

Supp. Table 9 displays the partial correlation analyses of model-based variables between the antisaccade and Simon task in Exp. 1. Supp. Table 10 displays the same analysis for purely behavioral variables. We applied Holm’s method to correct for multiple comparison. Supp. Fig. 2 displays the correlation between the ER on incongruent trials in the antisaccade and Simon tasks.

**Supplementary Table 9.** Correlation between parameter estimates across tasks in Exp. 1. The reaction time of controlled incongruent responses was significantly correlated. No parameter pertaining to inhibitory controlled was correlated across tasks. Multiple comparison correction by Holm’s method.

|  | Partial correlation<br>coefficient | P | P corrected |
| --- | --- | --- | --- |
| Automatic congruent RT | 0.029 | 0.610 | 0.610 |
| Automatic congruent<br>probability | 0.077 | 0.184 | 0.368 |
| Congruent controlled RT | 0.146 | 0.011 | 0.045 |
| Incongruent controlled RT | 0.312 | < 10<sup>-5</sup> | < 10<sup>-5</sup> |
| Congruent controlled<br>probability | 0.116 | 0.044 | 0.132 |

**Supplementary Table 10.** Correlation between behavioral outcomes in Experiment 1. Multiple comparison correction by Holm’s method.

|  | Partial correlation<br>coefficient | P | P corrected |
| --- | --- | --- | --- |
| Error rate incongruent<br>trials | 0.208 | < 10<sup>-3</sup> | < 10<sup>-3</sup> |
| Error rate congruent trials | 0.081 | 0.158 | 0.158 |
| Reaction time incongruent<br>trials | 0.309 | < 10<sup>-3</sup> | < 10<sup>-3</sup> |
| Reaction time congruent<br>trials | 0.207 | < 10<sup>-3</sup> | < 10<sup>-3</sup> |

**Supplementary Figure 2:**
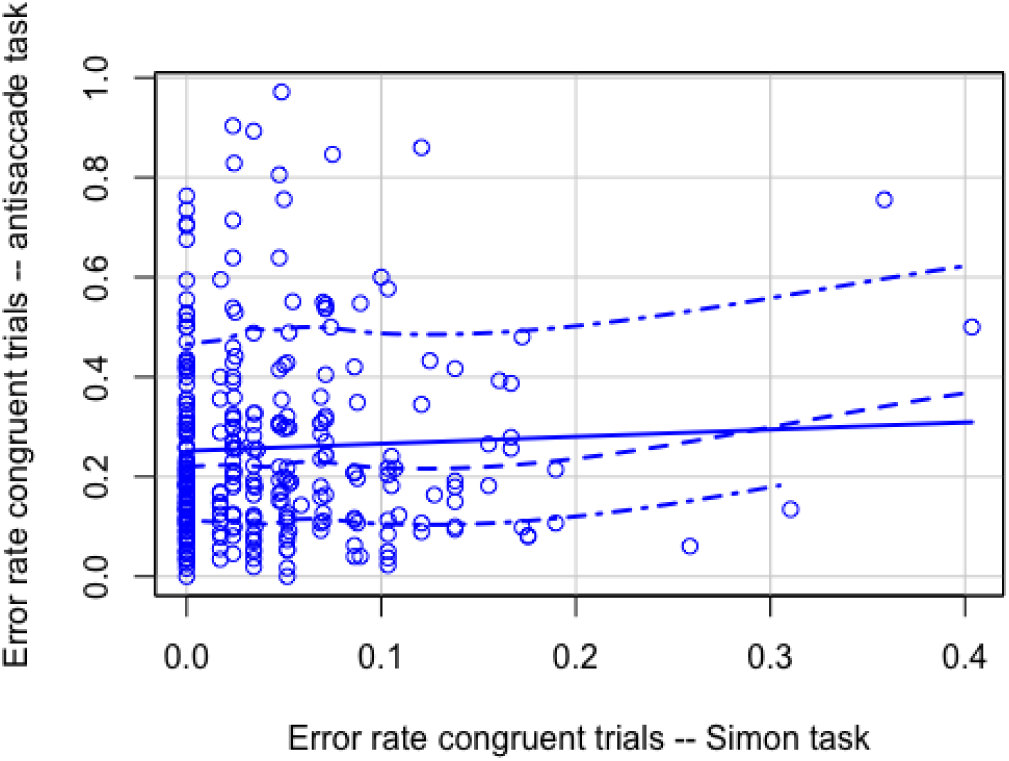
Correlation between the error rates on incongruent trials in the antisaccade and Simon tasks in Exp. 1 Circles represent individual participants. Blue solid line represents the regression slope, and the dotted line the running average of individual participants with its confidence interval displayed as dot-dash lines.

We performed a similar analysis in Exp. 2 (see Supp. Table 11 and 12) except that correlations were computed from estimates from a hierarchical SERIA model that controlled for i) conflict in the previous trial, ii) drug dose, and iii) subject entered as random effect. The subject specific intercepts (random effects) were used as random variables in the correlation analysis, because these represent the mean value of each subject after removing all other confounds.

### Supplementary material 4

**Supplementary Table 11.** Correlation between model-based variables across tasks in Exp. 2. There was a significant correlation between the reaction time of controlled responses across tasks. Multiple comparison correction by Holm’s method.

|  | Partial correlation coefficient | P | P corrected |
| --- | --- | --- | --- |
| Automatic congruent RT | 0.367 | 0.025 | 0.075 |
| Automatic congruent probability | 0.028 | 0.864 | 0.864 |
| Congruent controlled RT | 0.470 | 0.003 | 0.013 |
| Incongruent controlled RT | 0.478 | 0.002 | 0.013 |
| Congruent controlled probability | 0.277 | 0.095 | 0.191 |

**Supplementary Table 12.** Correlation between behavioral outcomes in Exp. 2. Multiple comparison correction by Holm’s method.

|  | Partial correlation coefficient | P | P corrected |
| --- | --- | --- | --- |
| Error rate incongruent trials | 0.422 | 0.009 | 0.027 |
| Error rate congruent trials | 0.314 | 0.058 | 0.065 |
| Reaction time incongruent trials | 0.457 | 0.004 | 0.017 |
| Reaction time congruent trials | 0.352 | 0.032 | 0.065 |

**Supplementary Figure 3:**
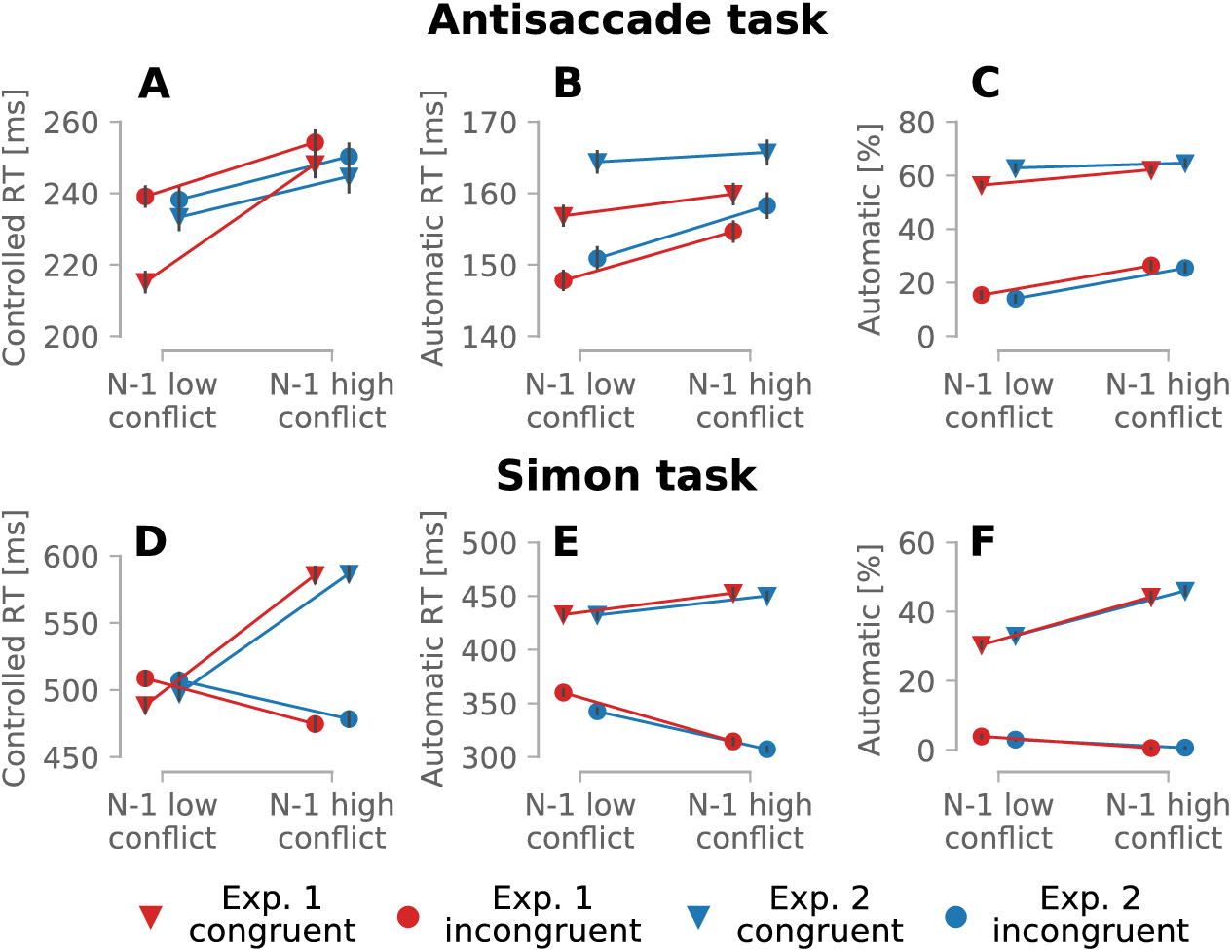
Model estimates of controlled and automatic RT as well as the percentage of automatic in Exp 1. and 2. A) Controlled RT in the antisaccade task, B) Automatic RT in the antisaccade task, C) Percentage of automatic responses in the antisaccade task, D-F) Similar to A-C in the Simon task. All parameter findings were replicated across experiments.

**Supplementary Table 13.** Mean model parameters as a function of drug. P values of the marginal effect of lorazepam.

|  | **Antisaccade task** |  |  |  |
| --- | --- | --- | --- | --- |
|  | Placebo | 0.5mg | 1.0mg | P |
| Percent automatic responses | 41 | 40 | 42 | 0.28 |
| Automatic RT [ms] | 155 | 159 | 163 | $< 10^{-5}$ |
| Controlled congruent RT [ms] | 230 | 240 | 246 | 0.012 |
| Controlled incongruent RT [ms] | 237 | 245 | 249 | 0.001 |
|  | **Simon task** |  |  |  |
|  | Placebo | 0.5mg | 1.0mg | P |
| Percent automatic responses | 19 | 23 | 19 | $< 10^{-5}$ |
| Automatic RT [ms] | 373 | 382 | 392 | $< 10^{-5}$ |
| Controlled congruent RT [ms] | 509 | 555 | 561 | $< 10^{-5}$ |
| Controlled incongruent RT [ms] | 472 | 493 | 511 | $< 10^{-5}$ |

## Notes

### Competing Interest Statement

I have read the journal's policy and the authors of this manuscript have the following competing interests: Eduardo A Aponte is a full time employee of Hoffman-LaRoche AG. All other authors have declared that no competing interests exist.

